# Tracking neural markers of template formation and implementation in attentional inhibition under different distractor consistency

**DOI:** 10.1101/2021.07.09.451861

**Authors:** Wen Wen, Zhibang Huang, Yin Hou, Sheng Li

## Abstract

Performing visual search tasks requires optimal attention deployment to promote targets and inhibit distractors. Rejection templates based on the distractor’s feature can be built to constrain the search process. We measured electroencephalography (EEG) of human participants of both sexes when they performed a visual search task in conditions where the distractor cues were constant within a block (fixed-cueing) or changed on a trial-by-trial basis (varied-cueing). In the fixed-cueing condition, sustained decoding of the cued colors could be achieved during the retention interval and the participants with higher decoding accuracy showed larger suppression benefit of the distractor cueing in the search period. In the varied-cueing condition, the cued color could only be transiently decoded after its onset and the higher decoding accuracy was observed from the participants who demonstrated lower suppression benefit. The differential neural representations of the to-be-ignored color in the two cueing conditions as well as their reverse associations with behavioral performance implied that rejection templates were formed in the fixed-cueing condition but not in the varied-cueing condition. Additionally, we observed stronger posterior alpha lateralization and mid-frontal theta/beta power during the retention interval of the varied-cueing condition, indicating the cognitive costs in template formation caused by the trialwise change of distractor colors. Taken together, our findings revealed the neural markers associated with the critical roles of distractor consistency in linking template formation to successful inhibition.

## Introduction

Human beings can allocate limited attentional resources to task-relevant items whilst filtering out irrelevant distractors. Extensive evidence has demonstrated that target templates held in memory facilitate visual performance (Desimone & Duncan, 1995; Eimer, 2014; Reeder & Peelen, 2013; Soto et al., 2005, 2008). By contrast, divergent findings have been reported regarding whether foreknowledge of the to-be-ignored feature, referred to as rejection templates (Arita et al., 2012; Woodman & Luck, 2007), can benefit search efficiency. Studies showed that the behavioral benefit of the rejection template is dependent on many factors (Beck et al., 2018; Conci et al., 2019; Han & Kim, 2009; Olivers, 2009; Tanda & Kawahara, 2019; Töllner et al., 2015; Zhang et al., 2020). Notably, the consistency of the to-be-ignored feature across trials was particularly prepotent (Cunningham & Egeth, 2016; Noonan et al., 2016; Vatterott & Vecera, 2012; Wen et al., 2018). These studies have demonstrated robust inhibition benefits when distractor cues remained unchanged within a block (fixed-cueing condition) and minimum benefits if not absent when the cues changed on a trial-by-trial basis (varied-cueing condition).

Researchers proposed that a stable rejection template is essential in distractor inhibition and can be created only under the fixed-cueing condition, whereas the varied-cueing context obstructed its formation (Arita et al., 2012; Carlisle, 2019; Cunningham & Egeth, 2016). So far, there was limited neural evidence shedding light on the existence of rejection templates. Reeder and colleagues (2017) found that negative cues that indicated the distractor feature led to decreased responses in the early visual cortex (V1-V3) prior to the search array. However, they later reported that this decrement was not a feature-specific suppression and there were no distinctive representations of the rejection templates (Reeder et al., 2018). Therefore, it is necessary to examine whether selective representations of the rejection templates could be identified.

Recent studies have reached the consensus that distractor inhibition could be learned when statistical regularities of distractors’ feature (Stilwell et al., 2019) or spatial location (Ferrante et al., 2018; Leber et al., 2016; Wang & Theeuwes, 2018) are expected (Geng et al., 2019). For example, Noonan et al. (2016) found decreased P1 component in the blocked condition where the distractor’s location was fixed. Similarly, van Moorselaar and Slagter (2019) showed that distractor-location learning induced behavioral benefits appeared after several repetitions and the learned expectation of distractors reduced stimulus-specific processing as reflected by the decreased Pd component and location-decoding accuracy. With all the evidence showing the influence of distractor consistency on stimulus processing, yet it remained less understood how distractor consistency modulates the process of template formation and subsequent implementation.

Mid-frontal theta-band activities have been linked to goal-related prioritization in working memory tasks (de Vries et al., 2020; Riddle et al., 2020) and conflict solving in the framework of cognitive control (Cavanagh & Frank, 2014; Cohen, 2014). Specifically, stronger theta activity was evoked by distractor cues relative to target cues during the delay period as a means of reducing interference from matched distractors (de Vries et al., 2019). Meanwhile, Dugué and colleagues (2014) observed stronger post-stimulus theta activity in correct trials than incorrect trials, revealing the association between post-stimulus theta power and target selection. Although behavioral studies have indicated the tight connection between template-based distractor inhibition and cognitive control (Han & Kim, 2009; Kiyonaga et al., 2012), the exact functional roles of theta activity in distractor inhibition, especially pertaining to template formation and implementation, need more empirical investigations.

The current study aimed to reveal the feature-specific representation of rejection template and the associated neural dynamics. Participants were provided with distractor cues and were asked to perform a cued visual search task (Figure 1A). We recorded electroencephalography (EEG) signals and applied multivariate decoding to reveal the neural representations of the to-be-ignored colors. We also examined the oscillatory activities corresponding to attentional selection, memory transformation, and cognitive control in the template formation and implementation stages.

**Figure 1.**
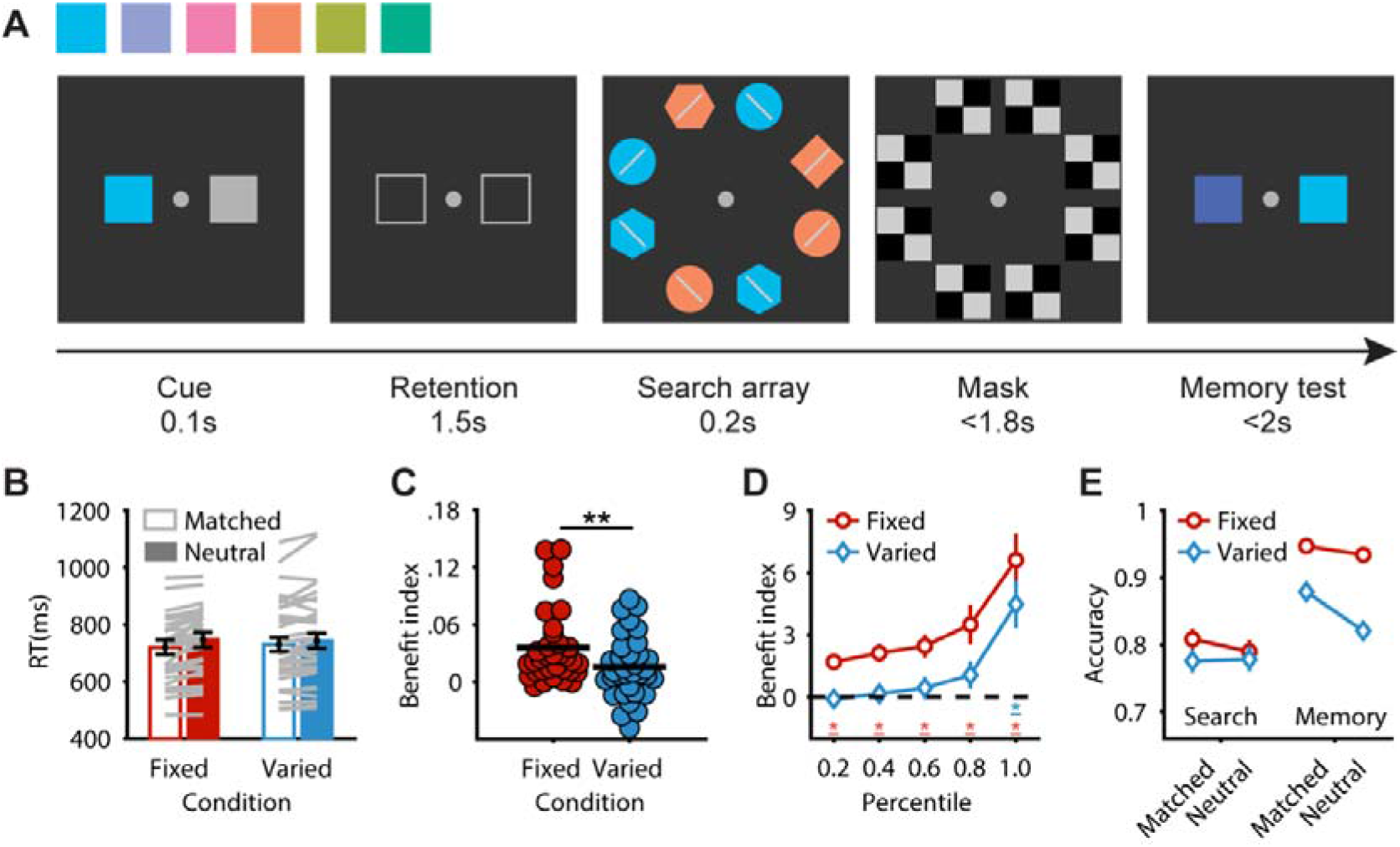
Task and behavioral results. (A) Task. The color cue indicated the to-be-ignored color in the search array. Participants were asked to find the target (diamond) and judge whether the bar inside of it was clockwise or counter-clockwise tilted relative to the mid-vertical. In the memory test, participants were instructed to select the color patch that matched the cue. (B) Search RT. Gray lines represent individuals’ data. Errorbars represent between-subject standard errors. (C) Suppression benefit index. Red and blue dots represent the individuals’ benefit indices in the fixed-cueing and varied-cueing conditions, respectively. Black lines represent the averaged benefit indices across participants. (D) Suppression benefit indices of RT percentile bins. Errorbars represent the standard deviations of 5000 times bootstrapped means. (E) Accuracy of visual search task and memory test. Errorbars represent the standard deviations of 5000 times bootstrapped means. * *p* < .05, ** *p* < .01, *** *p* < .001, 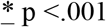 (two-tailed).

## Materials and Methods

### Participants

We tested thirty-four participants in total (mean age = 21.3, SD = 2.2), ranging from 18 to 27 years old, 15 females, 1 left-handed). According to the G-power estimation (Faul et al., 2007), thirty participants at least should be tested to guarantee a medium effect size (f = 0.25) and a high statistic power (90%). Participants were compensated with monetary rewards. All participants had normal or corrected-to-normal (color) vision and had no history of psychiatric or neurological disorders. All participants provided written informed consent prior to the experiment. The study was approved by the Committee for Protecting Human and Animal Subjects at the School of Psychological and Cognitive Sciences at Peking University (IRB protocol #2017-11-02).

### Stimuli and procedure

Participants were faced with a CRT monitor (refresh rate = 60 Hz) in a dark shielded EEG booth and their heads were stabilized using a chin rest. The schematic procedure of a trial is shown in Figure 1A. After a jittered blank (~1.5 s), a cue patch (1.5° × 1.5°) and an equal-sized gray patch were presented (1.5° to the center) for 0.1 s. The gray patch was presented to equalize the visual input in both hemispheres. The cue patch could appear at either hemifield with equal probability. Participants were instructed that target in the forthcoming search array would never be in the same color as the cued color. The color cue remained the same within one block in the fixed-cueing condition but changed on a trial-by-trial basis in the varied-cueing condition. Two gray frames of the same size and locations as the two patches were shown during the retention interval (1.5 s). The search array contained eight items of different shapes (diamond/circle/hexagon, in the same square size of 2°× 2°) and colors (four in target color and four in distractor color). Participants were instructed to find the target, defined solely by its shape (diamond), and discriminate the orientation of the bar inside of it. The target’s color was randomly selected in each trial. In matched trials (75%), the distractor’s color was the same as the cued color. In neutral trials (25%), the distractor’s color was a new color different from the target color and the cued color. The search array lasted for 0.2 s and was masked by checkerboards following its offset. Masks disappeared when participants made a response. A final memory test required participants to select the patch that was in the same color as the cue from two presented patches. To discourage participants from memorizing the cue color using semantic categorization, some pairs (~30%) in the memory test were set to be visually similar. Both the search task and the memory test had a response limit of 2 s.

Each cueing condition (fixed and varied) had 12 blocks with 72 trials per block. Six cue colors (selected from L*a*b color space, L = 70, equally distributed in the a*b plane) appeared in the first six blocks as well as the last six blocks in the fixed-cueing condition. Participants came twice and completed the fixed-cueing and varied-cueing conditions on separate days. The order of the conditions was counterbalanced between participants. Participants were instructed to look at the fixation dot and their eye movements were monitored using Eyelink 1000/1000Plus (1000 Hz, monocular).

### Statistical analysis

Only trials with correct responses in both the search task and the memory task were included for search task’s reaction times (RTs) analysis. Besides, we recursively removed outlier trials where search RTs were beyond 2.5 standard deviations from the mean. This resulted in 4.4% of the trials in the fixed-cueing condition and 3.7% of the trials in the varied-cueing condition being discarded. There was a general practice effect given that participants completed the two cueing conditions on separate days. Hence, we calculated the benefit index for each cueing condition to represent the RT facilitation resulted from the inhibition of matched distractors:

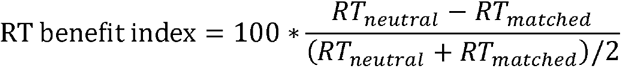

To examine the possible influence of testing order on the suppression benefit, we performed a two-way mixed ANOVA on benefit index with testing order (fixed-cueing first or varied-cueing first) as between-subject factor and cueing condition (fixed vs. varied) as within-subject factor. We further split the trials into five bins based on the search RT percentile for each trial type in each cueing condition. Then, we calculated the benefit index for each percentile bin in each cueing condition and conducted one-sample t-tests (two-tailed, alpha = .05) to compare the obtained benefit indices with zero. A two-way repeated ANOVA with factors of cueing condition (fixed vs. varied) and trial type (matched vs. neutral) was performed on the accuracies of the search task and the memory test. We reported the corrected degrees of freedom (Greenhouse-Geisser correction) when the equal variances assumption was violated.

### EEG recording

EEG data were acquired at a sampling rate of 1000 Hz using BrainAmps amplifier from a 64-channel EasyCap (Brain Products, Munich, Germany) with Ag/AgCl electrodes placed according to the 10-20 system. An external electrode placed below the right eye was used to measure eye movements. Electrode impedance was kept below 10 kΩ.

### EEG data analysis

#### EEG preprocessing

Offline preprocessing was performed with EEGLAB (Delorme & Makeig, 2004). After down-sampled to 500 Hz, the data was bandpass-filtered (1~30 Hz). Bad channels were spherically interpolated. We then re-referenced the data to the averaged signal of mastoids and performed independent component analysis (ICA) to correct ocular movements and other artifacts. On average, there were 3.8 (SD=1.4) and 4.1 (SD=1.5) components removed in the fixed-cueing and varied-cueing conditions respectively. Finally, the continuous EEG was epoched for further analyses (the exact epoch length and time-locked onset point were chosen depending on specific analyses). Besides behavioral exclusion described above, epochs in which any channel showed above-threshold amplitude (60 μV) at any time point of the epoch were removed. Since matched trials and neutral trials differed from each other only after the onset of search array, we used both types of trials when the analysis focused on the activities during the retention interval.

#### EEG decoding

To reveal the neural representations of the rejection templates, we performed color-decoding based on the whole-brain electrodes’ broad-band EEG data (EOG excluded) during the retention interval. The decoding was separately performed on cue-left and cue-right trials and the averaged results were used for statistical analysis. The continuous EEG was segmented into −0.2~1.6 s relative to the cue onset and baseline-corrected to the mean amplitude of the pre-stimulus period. We Gaussian-smoothed (window size = 16 ms) the data along the time dimension to reduce temporal noises. At each time point, data was partitioned into six training folds and one test fold. To create an unbiased classifier, the number of trials for each cue color in the training set was equalized by subsampling. Trial-averaged training sets were generated by averaging the trials from the same cue color. We first estimated the covariance matrix based on the trial-averaged training set. Next, we computed the pair-wise Mahalanobis distances between test trials and the trial-averaged training set using the covariance matrix (Wolff et al., 2017). A smaller distance toward the trial-averaged activation of one specific color cue condition indicated a larger pattern similarity. Therefore, the decoding was marked as a success when the distance between this given trial and the activation profile of the same color was smaller than any of other colors’ activation profiles in the training data. Cross-validation was carried out in a leave-one-fold-out manner. This routine was iterated 1000 times and trials were randomly assigned to the training and test set in each iteration. The averaged decoding accuracy was taken for the statistical analysis. The accuracy time series were Gaussian-smoothed (window size = 16 ms) and compared with the chance level (i.e., 1/6). Cluster-based permutation was performed on the decoding accuracy time-series over participants (alpha = .05, cluster-based nonparametric alpha = .05, cluster statistic = sum, two-tail, permutation times = 10000 (Maris & Oostenveld, 2007)).

#### Cross-temporal decoding

We followed the same decoding procedure as explained above except that we trained the model using data at one time point and tested it on all time points. We down-sampled the data into 50 Hz for computational efficiency. For the 2D result map (as in Figure 2B), the diagonal line represents the accuracies when training and testing were performed at the same time point while the off-diagonal points correspond to the temporal-generalized decoding. We adopted the leave-one-fold-out cross-validation method throughout the decoding process. Hence, the training-fold and the test-fold were never based on the same trials no matter whether the training and test time points were the same or different. This procedure was iterated 1000 times for each participant and the averaged result was used for group statistical tests. Cluster-based permutation test was conducted on the 2D result map at the participants’ population level (alpha = .01, cluster-based nonparametric alpha = .01, cluster statistic = sum, two-tail, permutation times = 10000 (Maris & Oostenveld, 2007)).

**Figure 2.**
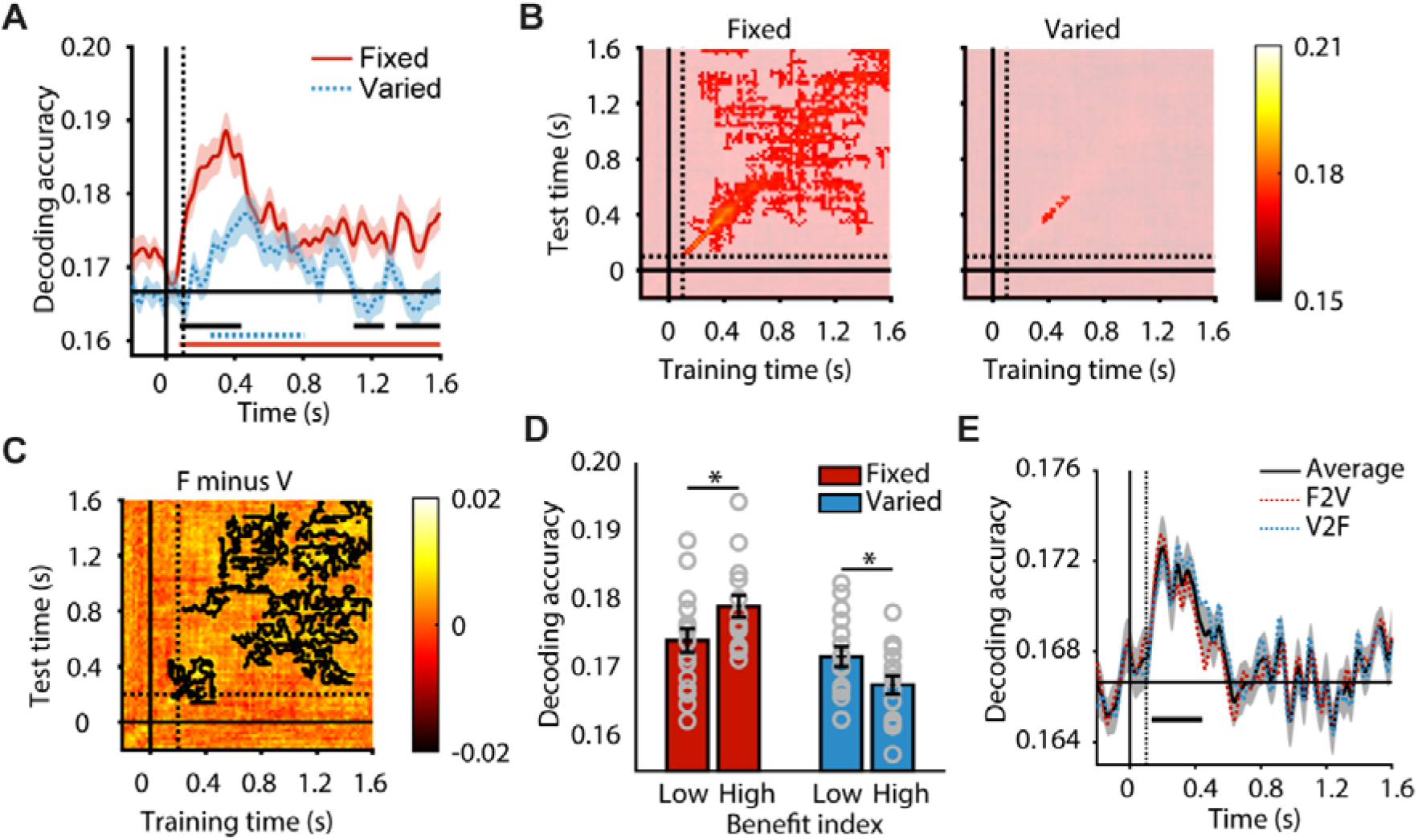
Results of color-decoding during retention interval. (A) Time-series of decoding accuracy in the fixed-cueing and varied-cueing conditions. The shaded areas represent the standard deviation of 5000 times bootstrapped mean. The solid/dashed vertical line indicate the cue onset/offset. Lines below the x-axis indicated significant time-clusters for each condition and the black lines correspond to the significant time-clusters for condition difference. (B) Cross-temporal decoding of cued color in the two cueing conditions. Alpha-blending was performed on insignificant time-points for better illustration. (C) Difference map of cross-temporal decoding. Black outlined areas indicate significantly higher decodability in the fixed-cueing condition compared to the varied-cueing condition as revealed by cluster-based permutation (alpha = .05, cluster-based nonparametric alpha = .05, cluster statistic = sum, two-tail, permutation times = 10000). (D) Decoding accuracies are shown for the low and high suppression benefit groups in both cueing conditions. Gray circles represent individual participants’ decoding accuracy, which is the average of the time-resolved decoding accuracies during retention interval (0~1600 ms). Errorbar represensts the between-subject standard error. \**p* <.05. (E) Cross-condition decoding. F2V stands for the decoding result when trained on data of the fixed-cueing condition and tested on data of the varied-cueing condition. The black time series is the average of bi-directional cross-condition decoding accuracy.

#### Cross-condition decoding

To compare the representations of color cues in the two cueing conditions, we performed cross-condition decoding by training the model using data from one condition and testing it on the other one (i.e., trained on the fixed-cueing condition and tested on the varied-cueing condition or vice versa). For each color of the training set, we subsampled equal number of trials to create an unbiased classifier. This subsampling process was iterated 1000 times. Bi-directional cross-condition decoding demonstrated similar results and therefore we performed statistical analysis on the average data (cluster-based permutation, alpha = .05, cluster-based nonparametric alpha = .05, cluster statistic = sum, two-tail, permutation times = 10000).

#### Multivariate topographical pattern similarity

The topographical activation pattern was defined as the whole-brain channels’ activities at a certain time point. First, we reorganized the data to integrate the left- and right-cued trials so that the left-hemifield electrodes were ipsilateral to the cue. Topographical pattern similarity was calculated as the correlation between the pattern of a given trial and the averaged pattern from the same cueing condition (i.e., within-condition scheme) or different cueing condition (i.e., between-condition scheme) (Figure 3A). The correlation was separately computed for each cued color to avoid the influence of sensory difference. For the between-condition scheme, the averaged topographical activation pattern was created by averaging trials of the same cued color from the other cueing condition. For the within-condition scheme, the averaged topographical activation pattern was created by averaging trials of the same cued color from the same cueing condition but in different blocks. We excluded the trials in the same block to reduce the temporal similarity effect. Considering that the number of trials that contributed to the averaged topographical activation patterns might confound the results, we subsampled the trials so that the number of trials used for the averaged topographical activation patterns across colors in each cueing condition was equalized. The subsampling process was iterated 1000 times. We treated the topographical activation pattern at each time point as a vector and calculated the Pearson’s correlation coefficients between the vector of the given trial and the vector of the averaged topographical activation pattern (Figure 3B). The averaged correlation coefficients across all interactions and all time points during the retention interval were used to represent each participant’s similarity scores in the four conditions (2 cueing conditions × 2 correlating schemes). The similarity scores were fisher-Z transformed when performing the statistical test at the population level.

**Figure 3.**
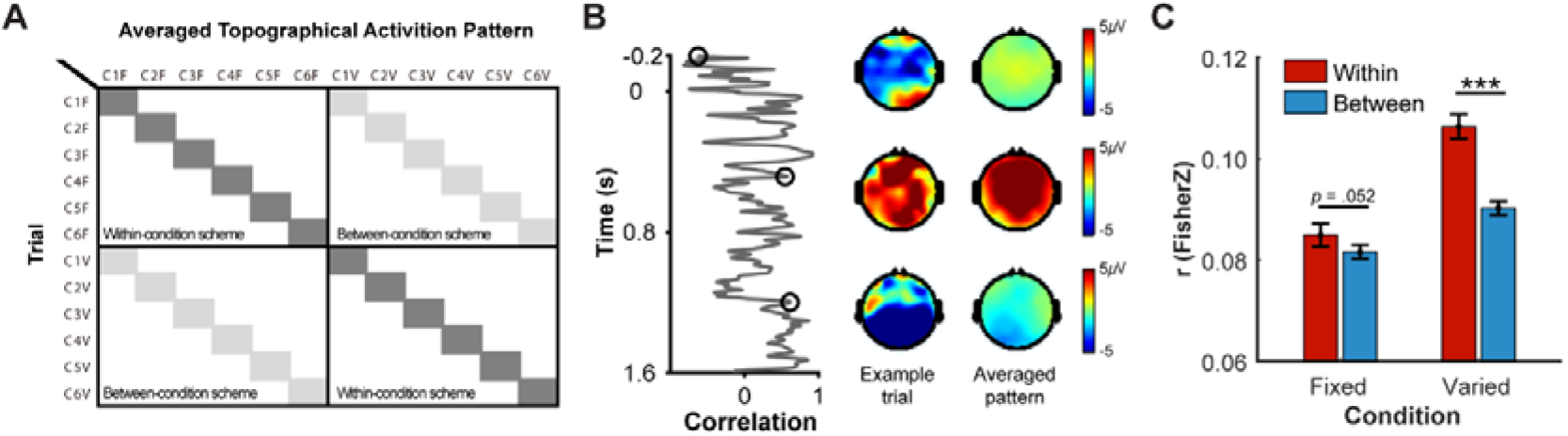
Multivariate topographical pattern similarity analysis. (A) Similarity matrix. C1F refers color1 in the fixed-cueing condition. Each row represents a given individual trial while each colume represents the averaged topographyical pattern of each color. Topographical similarities of highlighted filled grids were used for statistical analysis. (B) Schematic illustration of the topographical pattern similarity. The left panel is the correlation time series of the example trial and the averaged pattern during retention interval. The right side are the topographies of the example trial and the averaged pattern at three time points as indicated by the black circles in the time series. (C) Similarity scores. Errorbars correspond to the within-subject standard error.

#### Posterior alpha lateralization

A surface Laplacian filter (a 10th-order Legendre polynomial and a lambda of 10^-5^) was applied to increase topographical specificity and reduce the effects of volume conduction (Cohen, 2019). To calculate the posterior lateral alpha power, we segmented the continuous data into −0.4~1.6 s relative to the cue onset, and the averaged ERP was subtracted from each trial to remove phase-locked activity evoked by stimulus onset. Hilbert transform was used to extract the alpha-band (8~12 Hz) power of each trial using posterior occipital electrodes (P5/6, P7/8, PO7/8). Power data was decibel normalized to the averaged baseline (−0.4~−0.1 s). This process was separately performed on left-side cue and right-side cue trials and the averaged results were used for the statistical analysis. We calculated the alpha lateralization as the power difference between contralateral and ipsilateral electrodes (contralateral minus ipsilateral).

#### Time-frequency analysis

We performed time-frequency decomposition on the Laplacian-filtered data using Fieldtrip (Oostenveld et al., 2011). The continuous data were epoched −1.5~3 s relative to cue onset. After the time-frequency decomposition, we selected data between −0.4~1.6 s for further analysis. Each trial’s signal was decomposed using Morlet wavelet convolution every 50 ms. The frequencies ranged from 1~30 Hz in 30 linearly spaced steps and the width of the wavelet kernel increased linearly from 1 to 10 along with frequency bins. Normalization was done by decibel baselined to the pre-stimulus interval (−0.4~−0.1 s). Time-frequency decomposition was separately done on the left- and right-cued trials. We further combined the resulting data from the left- and right-cued trials so that the left-hemifield electrodes were ipsilateral to the stimulus. To test the difference between conditions, cluster-based permutation was performed at the group level. Adjacent time-frequency points exceeding the threshold (alpha = .01, two-tail) were grouped as a cluster. The cluster-level statistic was calculated by taking the sum of the difference values within the cluster (cluster alpha = .05). The number of random permutations using the Monte Carlo method was set to 10000.

We also examined the neural oscillations related to template implementation during visual search. Continuous data were epoched −1~2 s relative to search array onset. We adopted the same time-frequency decomposition method to obtain the oscillatory activities during −0.4~1.2 s for all trials.

## Results

### Behavioral suppression benefits in matched trials

As shown in Figure 1B, search RT had a significant main effect of trial type (F(1,33) = 21.132, *p* < .001, 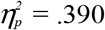) but not for cueing condition (F(1,33) = .016, *p* = .899, 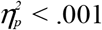), indicating that participants were generally faster in matched trials than neutral trials (19.8 ms). The significant interaction effect between cueing-condition and trial type (F(1,33) = 8.558, *p* = .006, 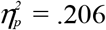) was driven by the larger suppression benefits in the fixed-cueing condition (26.8 ms, F(1,33) = 28.21, *p* < .001) than the varied-cueing condition (12.9 ms, F(1,33) = 7.17, *p* = .011). Considering the practice effect across testing days, we calculated benefit index which represents the suppression benefit of matched trials over neutral trials. Larger suppression benefits were observed in the fixed-cueing condition than the varied-cueing condition (F(1,32) = 9.661, *p* = .004, *η_p_^2^* = .232, Figure 1C) and testing order had no significant influence on suppression benefit (F(1,32) = 1.558, *p* = .221, *η_p_^2^* = .046). There was no significant interaction effect between the two factors (F(1,32) = .064, *p* = .802, *η_p_^2^* = .002).

Figure 1D demonstrates the benefit index across five RT percentile bins. Suppression benefits were evident across all bins in the fixed-cueing condition (all *p* < .001). However, for the varied-cueing condition, we observed significant suppression benefit only in the percentile bin with longest RTs (*p* < .001). The early and robust behavioral benefits in the fixed-cueing condition indicated the engagement of proactive inhibition when the to-be-ignored color remained the same. By contrast, the late suppression benefit of the longest RT bin in the varied-cueing condition implies that participants adopted a reactive control which required sufficient time to come into effect. These results were consistent with our previous findings that cognitive control played a crucial role in distractor inhibition in the varied-cueing condition (Wen et al., 2018).

We further analyzed the results of accuracy for visual search and memory test (Figure 1E). For the visual search accuracy, there were no significant differences between conditions (F(1,33) = 1.784, *p* = .191, *η_p_^2^* = .051) or trial types (F(1,33) = 3.115, *p* = .087, *η_p_^2^* = .086) but their interaction effect was significant (F(1,33) = 8.983, *p* = .005, *η_p_^2^* = .214). Simple effect analysis showed that participants were more accurate in the matched trials than the neutral trials in the fixed-cueing condition (F(1,33) = 11.31, *p* = .002). However, this difference was not evident in the varied-cueing condition (F(1,33) = 0.19, *p* = .662). For the memory test, participants were more accurate in the fixed-cueing condition (F(1, 33) = 100.318, *p* < .001, *η_p_^2^* = .752) and the matched trials (F(1, 33) = 93.234, *p* < .001, *η_p_^2^* = .739). The significant interaction effect (F(1, 33) = 46.202,*p* < .001, *η_p_^2^* = .583) indicated that when the cued color reappeared in the search array as the distractor color, it promoted memory performance to a larger extent in the varied-cueing condition (5.8%, F(1, 33) = 83.26, *p* < .001) than in the fixed-cueing condition (1.3%, F(1, 33) = 19.28, *p* < .001). This result might be explained by the refreshment strategy (or perceptual sampling) that using the reappeared matched distractors to enhance the memory of the to-be-ignored distractor color in the varied-cueing condition (Woodman & Luck, 2007).

### EEG decoding of the to-be-ignored color during retention interval

We performed multivariate decoding analysis on EEG data to reveal the representation of the distractor color during the retention interval. Color information could be decoded in both conditions (Figure 2A, fixed-cueing: 0.078~1.6 s, cluster *p* < .001; varied-cueing: 0.26~0.806 s, cluster *p* < .001). Importantly, the decoding accuracy was higher in the fixed-cueing condition compared with the varied-cueing condition, especially after the cue onset (cluster *p* < .001, 0.08~0.438 s) and prior to search array onset (cluster *p* = .021, 1.096~1.272 s; cluster *p* = .005, 1.342~1.6 s). To further rule out the possibility that the selective color representation was driven by the memory test, we split the participants based on their overall memory performance (across matched and neutral trials) and compared the decoding accuracy of the two subgroups in each cueing condition. No significant difference in decoding accuracy between the high and low performer groups was observed (fixed-cueing: t(32) = 0.433, *p* = .668; varied-cueing: t(32) = 0.155, *p* = .878).

Next, we focused on representation stability and performed cross-temporal decoding separately for the two cueing conditions. Figure 2B shows the two-dimensional maps of decoding accuracies. There was a wide range of generalization for color decoding between time points in the fixed-cueing condition, indicating a more stable representation of the cued color during the retention interval. In contrast, significant decoding performance was observed only when training and testing were performed on the data from the same time points in the varied-cueing condition, suggesting a highly dynamic representation of the cued color. Moreover, the difference map demonstrated that the fixed-cueing condition had higher off-diagonal decoding accuracy than the varied-cueing condition (Figure 2C). These results were in line with the larger behavioral suppression benefit in the fixed-cueing condition as such benefit would highly rely on the stable representation of the distractor’s feature to form an effective rejection template.

### Opposite effects in relating EEG decoding to behavior in two cueing conditions

To examine how the representations of distractor cues affect distractor filtering during visual search, we half-split the participants into two groups based on their behavioral benefit indices in each cueing condition. In the fixed-cueing condition, participants who showed larger suppression benefit had higher decoding accuracy (Figure 2D, t(32) = 2.119, *p* = .042, Cohen’d = 0.73). This pattern was reversed in the varied-cueing condition as higher decoding accuracy was observed among the low suppression benefit participants (Figure 2D, t(32) = 2.092, *p* = .044, Cohen’d = 0.71). This difference suggested that distinct neural representations of the to-be-ignored cue in the two cueing contingencies. To further examine the neural representation difference, we performed cross-condition decoding. Classifiers trained on trials from one cueing condition can only decode distractor cues from the other condition during the early retention interval (Figure 2E, 134~442 ms, cluster *p* < .001). The decoding accuracy was around chance level thereafter. This result indicated that the representation of color cues underwent two stages where sensory signals were emphasized in the initial stage. The unsuccessful cross-condition decoding suggested that different coding patterns were adopted to represent the color cue.

Additionally, we conducted multivariate topographical pattern similarity analysis on the basis of the within- and between-condition scheme correlations (Figure 3). If the rejection template was represented similarly in the two cueing conditions, one would expect that similarity scores for the within-condition and between-condition schemes were of no significant difference. However, the results revealed that the similarity scores were significantly different between the two cueing conditions (F(1,33) = 19.305, *p* < . 001, *ηp^2^* = 0.369) and the two correlating schemes (F(1,33) = 54.373, *p* < .001, *ηp* = 0.622). There was also an interaction effect (F(1,33) = 26.9, *p* < .001, *ηp^2^* = 0.449). Simple effect analysis revealed that the similarity score of the within-condition scheme was higher than that of the between-condition scheme for the varied-cueing condition (F(1,33) = 68.78, *p* < .001). The same trend was observed for the fixed-cueing condition with marginal significance (F(1,33) = 4.05, *p* = .052). These results suggest that, although exposed to the same sensory information, the brain signals during the retention interval manifested different topographical representations under the two cueing conditions.

### Posterior alpha lateralization during the early stage of retention interval

An increase in posterior alpha-band (8~12 Hz) activity has been regarded as a neural marker of attentional inhibition (Jensen & Mazaheri, 2010; Kelly et al., 2006; Worden et al., 2000). Correspondingly, the decrease of alpha power at the contralateral side of an attended stimulus indicated the release from inhibition (i.e., increase in attention). As shown in Figure 4A, we observed significant alpha lateralization (contralateral minus ipsilateral) induced by the color cue in both cueing conditions (fixed-cueing condition: 0.282~0.74 s, cluster *p* = .008; varied-cueing condition: 0.266~1.124 s, cluster *p* = .001). Stronger alpha lateralization (herein, more negative) was found in the varied-cueing condition relative to the fixed-cueing condition (0.338~1.066 s, cluster *p* = .003), indicating a larger attentional bias towards the color cue at the initial stage of the retention interval. This attentional bias was attenuated with the repetition of color cues in the fixed-cueing condition.

**Figure 4.**
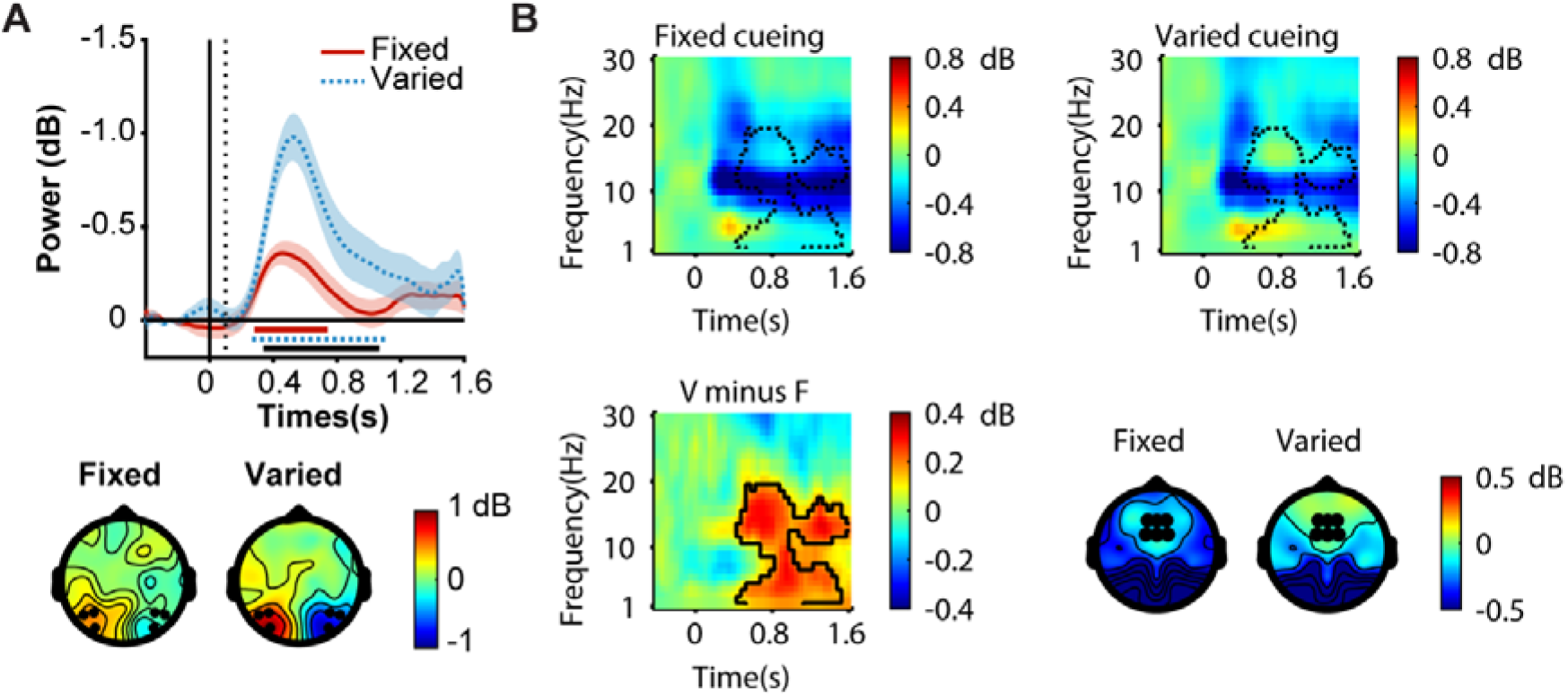
Oscillatory activities during retention interval. (A) Top: Posterior lateralized alpha power in the fixed-cueing and varied-cueing conditions. The shaded areas represent the between-subject standard error. The solid/dashed vertical line indicated the cue onset/offset. Lines below the x-axis indicated significant time-clusters for each condition with the black line corresponding to their difference. Bottom: Topography of the averaged lateralized alpha power during 300~800 ms. Topographical maps indicate the difference values of alpha power between left-cue minus right-cue trials. Since contralateral alpha power would decrease relative to ipsilateral alpha power when distractor cues are attended, left posterior electrodes are positive in this plot. The black dots highlight the channels used for alpha lateralization analysis (P5/6, P7/8, PO7/8). (B) Time-frequency map extracted from mid-frontal electrodes (FCz, FC1, FC2, Fz, F1, F2). Dotted contours were adopted from the time-frequency difference map (varied minus fixed). Significant time-frequency clusters from the non-parametric permutation test are outlined by the dark contour. Data was extracted from mid-frontal electrodes. Right-bottom: Topographical distribution of mid-frontal theta activity (3~7 Hz, 0.1~1.6 s) for the fixed-cueing and varied-cueing conditions. Black dots correspond to the channels used for time-frequency analysis.

### Mid-frontal theta and beta activities during the late stage of retention interval

Next, we focused on mid-frontal activities in the two cueing conditions (Figure 4B). The time-frequency difference map of mid-frontal electrodes showed that varied cues induced stronger oscillatory activities in the range of theta-band and beta-band at the later stage of the retention interval in comparison with the fixed cues (0.35~1.6 s, 2~19 Hz, cluster *p* = .002). Frontal theta oscillation (3~7 Hz) has been regarded as a reflection of top-down control in goal-directed cognitive processes (Cavanagh & Frank, 2014). In two recent studies, stronger mid-frontal theta was observed during the retention interval if the cue indicated the distractor rather than the target (de Vries et al., 2019) or switching rather than repeating the task sets (Cooper et al., 2019). The enhanced mid-frontal theta activity was also suggested to be associated with the prevention of interference from the anticipated distractor (de Vries et al., 2019) or the proactive control of task set switching (Cooper et al., 2019). In our experiment, the varied-cueing condition shared these critical aspects of task structure with these two studies. Therefore, we consider that both the prevention of interference from the anticipated distractor and the proactive control of task set switching could contribute to the observed mid-frontal theta effects during the retention interval.

On the other hand, frontal beta activity (13~30 Hz) has been proposed to reflect a short-lived transition period between representations (e.g., from latent to active) late in the WM delay in the service of task demands (Spitzer & Haegens, 2017). We speculated that the higher mid-frontal beta activity in the varied-cueing condition (Figure 4B) reflected a stronger engagement in transferring the cueing information from perceptual representation into rejection template that can be used to filter out the matched distractors in the following search task. However, given the lack of literature on the roles of frontal beta in attentional suppression, future experiments with more specific designs are required to test this interpretation.

### Mid-frontal theta activity during visual search period

Finally, time-frequency analysis also highlighted the dominant low-frequency oscillatory activity in mid-frontal electrodes during the search process (1~7 Hz, 0.2-1.2 s, cluster *p* < .040, Figure 5A). There were no significant differences between conditions, trial types, or their interaction (repeated measures ANOVA, all *ps* > .302). We then examined the functional relevance of the post-stimulus theta power with search performance (Figure 5B). Particularly, theta power predicted search RT in the varied-cueing condition (matched trials: Spearman’s ρ = −.564, *p* = .001; neutral trials: Spearman’s ρ = −.680, *p* < .001) but not in the fixed-cueing condition (matched trials: Spearman’s ρ = −.174, *p* = .324; neutral trials: Spearman’s ρ = −.108, *p* = .542). These results indicate that mid-frontal theta at the post-stimulus stage reflected a reactive control process that filtered out the distractors in the absence of stable rejection templates.

**Figure 5.**
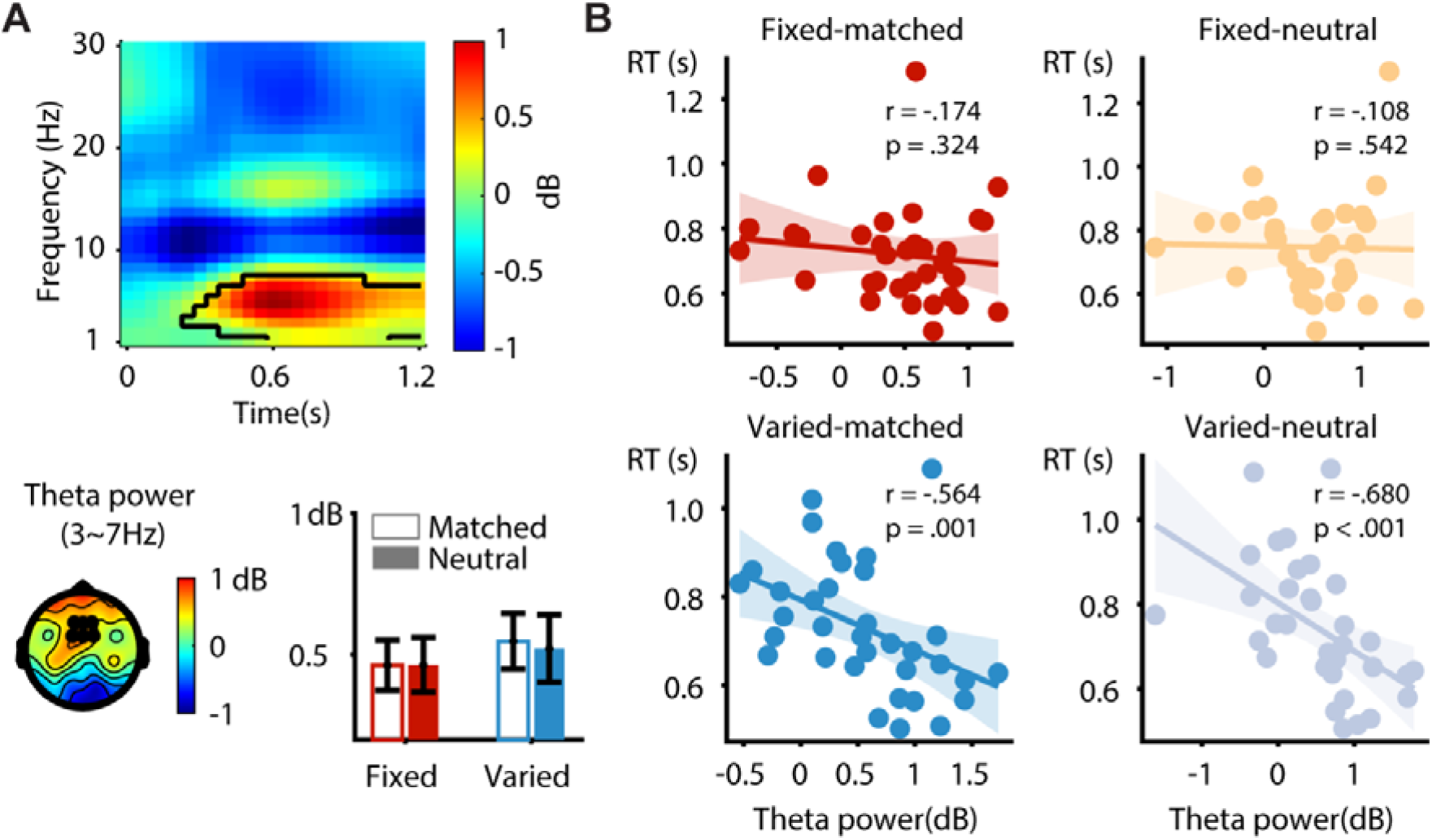
Mid-frontal theta after search array onset. (A) Top: Time-frequency map after search array onset. The significant time-frequency cluster from the non-parametric permutation test was outlined by the dark contour. Topography reflects averaged theta power during 0.2~0.7s. The time window was chosen to avoid stimulus-driven and response-related contaminates. The average search RTs across conditions was 735 ms. The black dots indicated the channels used for power analysis (FCz, FC1, FC2, Fz, F1, F2). Bar plot shows the average mid-frontal theta power within time window of interest. Errorbars represent the between-subject corrected standard error. (B) Spearman-rank correlations between frontal theta power and RTs in matched trials and neutral trials of two cueing conditions. The shaded area represents the 95% confidence interval for the fitted line.

## Discussion

Distractor inhibition is challenging and context-dependent. Inconsistent results have been reported concerning whether distractors matched with the to-be-ignored feature could be inhibited. The present study identified the neural signature of the rejection template in the fixed-cueing condition as characterized by the sustained decoding of the to-be-ignored color and the stable cross-temporal decoding performance. By contrast, trialwise updates of distractor cues in the varied-cueing condition hampered the establishment of stable rejection templates. This was reflected as the smaller behavioral suppression benefit and the transient decoding performance that did not persist in the late stage of the retention interval. Moreover, we found that better neural representation of the distractor cues in the varied-cueing condition led to smaller suppression benefits. These findings collectively suggested that creating rejection templates is difficult when the distractor consistency is violated.

The specific processes related to template formation were unfolded by oscillatory activities in different frequency bands. First, we observed stronger posterior alpha lateralization toward the color cue in the varied-cueing condition than the fixed-cueing condition, reflecting the higher necessity of paying attention to the color cue when it changed on a trial-by-trial basis. The posterior alpha lateralization was followed by mid-frontal theta and beta activities in the late stage of the retention interval. Based on previous literature, we suggest that the theta band activity reflected the control processes such as the prevention of the anticipated distractor interference and the proactive control of cue switching (Cooper et al., 2019; de Vries et al., 2019). The beta band activity, on the other hand, may index the memory transition process that transferred the color cue from perceptual representation to rejection template (Spitzer & Haegens, 2017). However, this tentative explanation needs future empirical investigations.

Capitalized on color decoding of EEG signal, we demonstrated selective representations of the to-be-ignored feature in both the fixed-cueing and varied-cueing conditions. In Reeder and colleagues’ (2018) fMRI study, they performed representational similarity analysis to calculate the distance metric of five cued colors in target, distractor, and neutral cueing conditions. Compared with the neutral condition, a more distinctive representation was observed only when the color was cued to be the target but not when the color was a distracting feature. The authors argued that the representations of general task-irrelevant features in neutral condition might not be an ideal baseline because participants might inhibit the neutral cue as well. Instead of comparing the representation distinctiveness across different cueing contingencies, we emphasized selective representation of the to-be-ignored feature within each condition. Meanwhile, the decoding accuracy time series provided us with a temporally fine-scaled picture of the representations of rejection templates. In another study, by repeating distractor locations, van Moorselaar and colleagues (2019) showed how learned expectations promoted distractor inhibition. Whereas distractor expectation reduced distractor-specific processing after search array onset, meaningful preparatory neural tuning to distractor locations was not observed in their study. Herein, we first provide the evidence of color-selective representation of the distractor cue during the retention interval, particularly in the fixed-cueing condition. Discrepancies in the contents of rejection template as well as the experimental design might contribute to the different findings. Compared to the location-based expectation in van Moorselaar’s study which only excluded one item, taking the advantage of the to-be-ignored color in the present study would help to filter out half of the items in a search array. Moreover, fixed cues that were presented within one block could further facilitate the generation of stable rejection templates.

The constant cueing contingency in fixed-cueing condition promoted the expectation of distractor color and therefore boosted the color selectivity during the retention interval. This might account for the very early (<100 ms) emergence of the template information extracted by decoding. In fact, previous work showed that expectation could pre-activate representations of expected sensory stimuli (Kok et al., 2017) and strategically bias gazes toward the foreknown distractor in the preparatory stage to reduce stimulus-evoked attractions (Wen et al., 2021). Behaviorally, fixed cues also facilitate distractor suppression which echoed previous findings that participants can be more effective in excluding the irrelevant items in a predictive context (Cunningham & Egeth, 2016; Noonan et al., 2016; Töllner et al., 2015).

The opposite effects in relating decoding accuracy to suppression benefit in the two cueing conditions suggest that neural representations of the distractor cues were different in nature. Together with oscillatory activity in retention, this reverse pattern implied that the formation of rejection template was dependent on distractor consistency. Since robust rejection templates were built in the fixed-cueing condition, participants could implement the rejection template in this condition to proactively suppress the matched distractors, but relied on reactive control to filter out non-targets in the varied-cueing condition. This was evidenced by the negative correlation between the post-stimulus theta power and search RT that was only observed in the varied-cueing condition. In addition, behavioral suppression benefits in the varied-cueing condition were smaller than the fixed-cueing condition and appeared only in the longest RT bin. Hence, while proactive inhibition played a central role in the fixed-cueing condition, distractor suppression in the varied-cueing condition was primarily undertaken by reactive control. Notably, the negative correlation was observed in both matched trials and neutral trials, implying that reactive control operated in a universal manner and rejected all nontargets in the absence of a stable rejection template.

Given the current design, we could not elucidate how the matched distractors were filtered out based on the rejection template especially in the fixed-cueing condition. Previous work found that the target template was pre-activated briefly before stimulus onset and served as a match filter to convert the sensory input of stimulus into the decision-relevant representation (Myers et al., 2015). However, it remains unknown whether the rejection template functions the same way as the target template. It has been proposed that the rejection template might reduce the activation of matched features and therefore prevent those matched distractors from interfering with target processing in the first place (Moher et al., 2014; Noonan et al., 2016; van Zoest et al., 2021). Spatial tuning to the distractor location also decreased when participants learned to ignore the distractor (van Moorselaar & Slagter, 2019). Our behavioral results showed that inhibition was achieved even in the fastest RT bin in the fixed-cueing condition, indicating a proactive suppression presumably through the reduction of the distractor-evoked response. On the contrary, rejection templates could be utilized reactively (Moher & Egeth, 2012) and contribute to suppression benefits of the longest RT bin in the varied-cueing condition. Future studies are required to further investigate the factors that modulate the implementation process.

In conclusion, our results revealed the neural markers for the distinctive processes of template formation and implementation between the fixed-cueing and varied-cueing conditions. Distractor consistency in the fixed-cueing condition promotes the formation of rejection template and leads to the significant behavioral suppression benefit. In the varied-cueing condition, transient decoding performance as well as the stronger posterior alpha lateralization and mid-frontal theta/beta power during the retention interval imply the cognitive costs in template formation caused by the trial-by-trial changes of the distractor features. In comparison to the fixed-cueing condition where rejection template can be implemented to proactively suppress matched distractors, the varied-cueing condition relies on reactive control to filter out non-targets. Together, these findings provide an integrative view of template-guided distractor inhibition in visual search.

## Acknowledgments

This work was supported by a grant from National Key R&D Program of China (2017YFB1002503).

